# Asymmetric increase in episodic and procedural memory interference in older adults

**DOI:** 10.1101/2025.06.19.660650

**Authors:** Busra Celik, Lara Kamal, Dora Goldstein, Griffin Alpert, Samantha Gonzalez, Vian Nguyen, Michael Freedberg

## Abstract

In younger adults, newly formed procedural memories are weakened by the subsequent formation of episodic memories (E→P interference) and vice versa (P→E interference; “cross-memory interference”). Older adults experience significant decline in episodic memory but maintain relatively intact procedural memory. This asymmetric decline in memory may also cause an asymmetric change in cross-memory interference compared to younger adults. For example, older adults may experience a significant increase in one type of cross-memory interference while leaving the other unchanged. Additionally, decline in episodic memory may cause E→P interference to either increase or decrease depending on how the episodic and procedural memory systems interact. However, no study to our knowledge has compared cross-memory interference between younger and older adults. We investigated cross-memory interference in younger and older adults by measuring E→P (Exp. 1) and P→E (Exp. 2) interference in 40 younger (18-40 years old) and 40 older (≥ 55 years old) adults. Compared to younger adults, the results show that older adults experience significantly stronger E→P interference while P→E interference was statistically indistinguishable between groups. These results confirm that older adults experience an asymmetric increase in cross-memory interference and suggest that the increase in E→P interference is related to the asymmetric decline in episodic memory relative to procedural memory.

## Introduction

Human memories are often categorized as either episodic or procedural [1]. Episodic memories include verbalizable, autobiographical facts, events, and objects [2], and are supported by a distributed network of hippocampal-cortical connections [3,4]. Episodic memory is commonly studied using list-learning paradigms, such as the Rey Auditory Verbal Learning Test [5].

Procedural memories are cognitive and sensorimotor habits and skills gradually learned through practice [6,7] that are supported by striatal-cortical loops [8–10]. Procedural memory is commonly studied using the serial reaction time task (SRTT; [11]).

In younger adults, acquiring episodic memories immediately after learning a procedural skill weakens the wakeful consolidation of the procedural skill (E→P interference) and vice versa (P→E interference; [12–16]). These phenomena (hereafter referred to as “cross-memory interference”) are mechanistically linked to episodic and procedural memory network function. For example, stimulation-induced disruption of the motor cortex (part of the motor procedural network) reduces P→E interference and the same disruption of the dorsolateral prefrontal cortex (part of the episodic memory network) reduces E→P interference [15]. Thus, cross-memory interference is thought to result from interactions between episodic and procedural memory systems.

Age-related memory loss is a common concern for older adults [17–19]. However, aging does not uniformly affect memory [20,21]. While procedural memory is relatively spared (Brown et al., 2009), episodic memory declines substantially [20,21]. Episodic memory deficits can occur for a number of reasons, leaving the source of these deficits an open area of investigation. However, these declines are commonly reported to be associated with decreased engagement of the episodic memory network [22–25].

The asymmetric decline in memory in older adults raises an important question related to cross-memory interference: Do asymmetric changes in episodic and procedural memory also cause asymmetric changes in cross-memory interference? The answer to this question relies on a comprehensive understanding of how cross-memory interference occurs. For example, one can argue that general age-related changes in neural processes, such as global network dedifferentiation [26,27] could reasonably increase E→P and P→E interference. Alternatively, one can argue that cross-memory interference is specifically related to the health of each memory system and that decline in only one system would produce asymmetric changes in cross-memory interference. Furthermore, if these changes are asymmetric in that one type of interference becomes exacerbated more than the other, would decline in one type of memory compared to the other [20,21] increase or decrease interference? In the case of older adults, decline in the health of the episodic memory system could lead to decreased regulation of communication between the episodic memory system and other networks, which may increase E→P interference. Alternatively, one could argue that decline in the episodic memory system could reduce the strength of inhibitory signaling on the procedural memory system, thus reducing E→P interference.

Although the answers to these questions rely on a more nuanced mechanistic understanding of how cross-memory interference occurs, which is beyond the scope of this article, it is first necessary to establish whether cross-memory interference occurs in older adults and how it compares to younger adults. To this end, we compared P→E and E→P interference between cognitively unimpaired younger and older adults by performing a conceptual replication of Brown and Robertson’s (2007) original experiment, which showed that younger adults experience E→P (Experiment 1) and P→E (Experiment 2) interference. Here, we provide, to our knowledge, the first direct comparison of cross-memory interference between younger and older adults.

## Methods

### Study overview

We designed a conceptual replication of Brown and Robertson’s (2007) experiments, which revealed evidence of E→P and P→E interference [13] in younger adults. Participants in Brown and Robertson’s (2007) experiments visited the laboratory in the morning and returned twelve hours later. During the morning visit, participants either acquired episodic or procedural memories. After memory acquisition, half of the participants acquired memories of the other type, while the other half performed a control task. Twelve hours later, all participants were asked to recall the memories they acquired from the first task they performed in the morning. Consolidation was defined as the difference in memory performance between the morning and follow-up sessions.

The goals of the current study were to determine whether the magnitude of P→E and E→P are significantly different between cognitively unimpaired younger (18-40 years old) and older adults (≥55 years old). In the context of our study, interference occurs when the consolidation of one memory type is weakened by subsequent acquisition of the other.

Experiments 1 and 2 investigated P→E and E→P interference, respectively. The experiments consisted of morning and afternoon sessions. Immediately before the first experimental session, participants provided their informed consent to participate in the study, were screened for eligibility using the Mini-Mental Status Exam (MMSE; [28]), and completed the Everyday Memory Questionnaire (EMQ; [29]). The MMSE was used to exclude participants with a high possibility of dementia (score < 24) and the EMQ was used to explore whether self-perceived occurrence of memory failures in daily life is associated with episodic memory interference.

Each age group (younger and older) was divided into two sub-groups, which differed only in whether participants also performed a task to learn memories of the other type (see *Procedures*). Thus, each experiment included four groups: older adults-learning, older adults-control, younger adults-learning, and younger adults-control.

### Experiment 1 (E→P interference)

De-identified data (CSV files) and code (R markdown file; R-Studio version 4.2.3) needed to reproduce analyses in this experiment are available for download at OpenICPSR.org (https://www.openicpsr.org/openicpsr/project/208584/version/V2/view). The experiment was pre-registered on aspredicted.org (#137403) after 19 subjects’ data were collected but before our statistical analysis. We predicted that older adults would show significantly greater E→P memory interference than younger adults.

### Participants

Data collection for this experiment took place between January 1^st^, 2023, and June 1^st^, 2024. Forty participants from the greater Austin area provided their informed consent to participate and privacy rights of all participants were observed by all experimenters. Eligible participants were between the ages of 18-40 years (younger adults) or 55 years or above (older adults), scored ≥ 24 on the MMSE [28], had normal color vision (self-reported), and normal or corrected-to-normal vision. No participants met criteria for exclusion. Data were analyzed for 20 younger (14 Female; 24.64 ± 4.46 years, 6 Male; 23.66 ± 2.50 years) and 20 older (16 Female; 61.93 ± 5.96 years, 4 Male; 64.25 ± 7.27 years) participants. The Institutional Review Board at UT Austin approved all procedures on March 19^th^, 2021 (STUDY #00000860), and all participants were compensated for their participation. We determined our sample size based on the first experiment in Brown and Robertson’s (2007) report, which revealed a significant interfering effect of episodic memory acquisition on procedural memory consolidation in healthy younger adults: While the control group experienced offline gains between laboratory visits (n=10; Mean = 20, SEM = 7), the control group experienced offline losses (n=10; Mean = -22, SEM = 14).

This difference equates to an effect size of 1.2 (Cohen’s *d*). Assuming an alpha of 0.05 and power equal to 0.85, a sample size of ten participants per group would yield an 85.9% chance of observing a significant memory interference effect in our younger adult sample. Thus, all four groups in Experiment 1 included ten participants.

### Procedures

Figure 1 **(top)** shows the experimental timeline for Experiment 1. In the morning, participants performed the SRTT (P^1^). Next, depending on the sub-group, participants either performed an episodic learning (word list learning task) or control (vowel counting) task. Finally, after a 6-12 h interval (average = 6 hours and 35 minutes ± 30.97 minutes), participants were retested on the SRTT during the afternoon session (P^2^). After retesting, we measured participants’ conscious control of the SRTT sequence using the PDP [30]. The rationale for including this measurement is that conscious sequence awareness affects the consolidation of procedural memories (Robertson et al., 2004) and we wanted to provide statistical control of this variable in our analysis (see *Statistical Analysis*).

**Figure 1.**
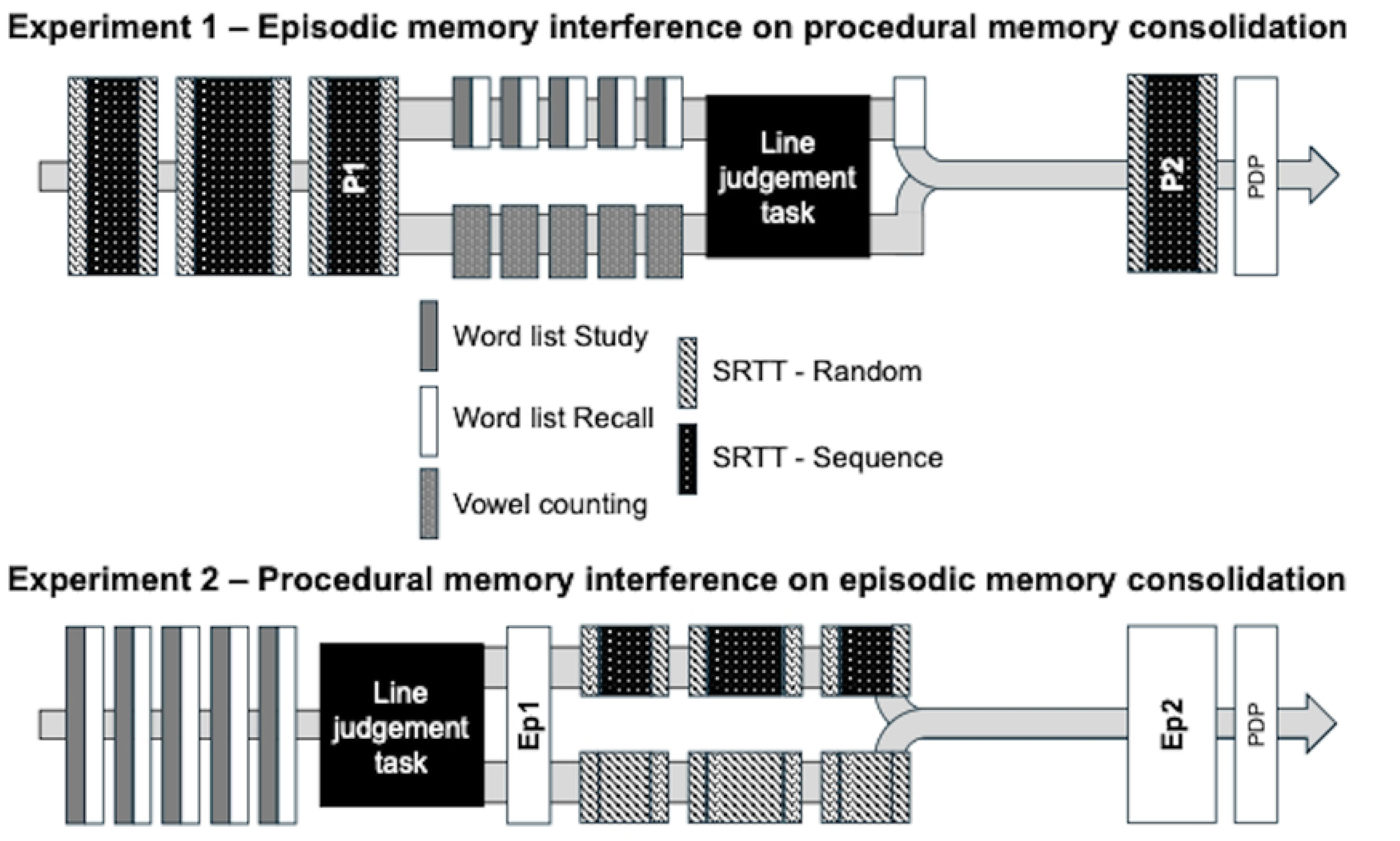
Experimental timeline for both experiments. Consolidation in Experiments 1 and 2 was calculated as P^2^ minus P^1^ and Ep^2^ minus Ep^1^, respectively.

#### SRTT (P^1^)

Participants were seated in front of a computer screen and placed the four non-thumb fingers of their right hand on a button box (Chronos Response Device, Psychology Software Tools), such that their index finger rested on the leftmost button (1), followed by their middle (2), ring (3), and pinky (4) fingers. Because the Chronos response box has five buttons, one button was occluded, and participants were instructed to ignore it. On the center of the computer screen, participants saw four white hollow circles arranged horizontally. On each trial, one of the hollow circles changed to a filled white circle, indicating that the participant needed to depress the corresponding response box button as quickly and accurately as possible.

Participants initially performed a practice block with random cues to achieve task familiarity and then performed three blocks of the actual task. The first, second, and third blocks contained 280, 400, and 280 trials, respectively. Each block included 50 random trials at the beginning and end to discourage the formation of explicit sequence awareness. The remaining trials followed a 12-element sequence (2-3-1-4-3-2-4-1-3-4-2-1). Thus, across the three blocks of the morning session, participants performed the sequence 15, 25, and 15 times.

#### Episodic learning task (word list learning)

Participants were instructed to memorize 16 words individually displayed on a computer screen: Age, Body, College, Door, Figure, Head, Keep, Level, Music, Night, Office, People, Room, Surface, Town, and Week. Once all the words were shown, participants’ were directed to recall the words in any order with no time limit imposed. This was repeated five times to encourage episodic memory acquisition. Next, participants performed a ten-minute perceptual distraction task (line judgment task). We included the line judgment task to prevent participants from verbally rehearsing the list of words between episodic memory acquisition and recall, thus minimizing interindividual and intergroup differences in the use of rehearsal as a cognitive strategy to increase episodic memory performance. Finally, participants performed a final memory recall trial without exposure to the list. Performance on this recall trial was used to measure participants’ episodic memory performance.

#### Episodic control task (vowel counting)

The episodic control task was similar to the episodic memory task but did not include instructions to memorize words. Participants were shown a string of randomly generated letters 3-12 letters long and were instructed to verbally indicate the number of vowels in each string. Like the episodic learning task, participants observed individual strings of letters for 16 trials per block for a total of 5 blocks (90 total trials) and this was followed by 10 minutes of the line judgment task.

#### SRTT (P^2^)

Participants re-performed the SRTT six to twelve hours later for a single block. This block was identical to Block 3 performed during P^1^ (Fig. 1**, top**).

#### Awareness Testing

The PDP is designed to assess conscious control of sequential information [30]. We included this measure to isolate statistical variance due to sequence awareness from our fixed effects of interest. The PDP includes two parts: Part 1 requires participants to complete 12 unique triplets of the sequence, while Part 2 requires participants to *avoid* the third element of the same triplets. During Part 1, participants are guided to press the first two elements of the sequence and then must indicate the third element with a single keypress.

During Part 2, participants perform the same task but must press either of the buttons that do not complete the triplet. Participants were not allowed to respond with the second element of the triplet, as no repetitions were allowed to occur during the performance blocks. Each part included 12 trials, corresponding to each unique triplet in the sequence (e.g., 2-3-1, 3-1-4, 1-4-3, etc.).

### Data Analysis

Response times (RTs) on the procedural task were defined as the duration to produce an accurate response on each trial. Any trial with a response time exceeding 2.7 standard deviations (the top 1% of RTs) from a participant’s mean RT was excluded from statistical analysis. Procedural memory consolidation was calculated as the difference in skill performance across P^1^ and P^2^ (Fig.1**, bottom**; P^2^ minus P^1^), where positive values indicate offline gains and negative values indicate offline losses. Procedural memory performance at P^1^ and P^2^ were calculated using the third block of P^1^ and the sole P^2^ block, respectively, as the average difference in RT between the final 50 random and sequenced trials [13]. SRTT accuracy scores are not a meaningful measure of skill performance since error frequencies are exceedingly minimal [31,32]. Therefore, we did not analyze these data. Performance on the episodic list-learning task was calculated as the number of recalled words on the final recall trial. To obtain awareness scores, we calculated the proportion of correctly performed trials for Parts 1 (sequence inclusion) and 2 (sequence exclusion) of the PDP. Then, we subtracted 0.33 from Part 1 scores and 0.67 from Part 2 scores, which correspond to chance performance. Finally, we summed the differences to create a composite awareness score for each participant that ranged from -1 (below chance performance) to 1 (above chance performance).

### Statistical Analysis

Our statistical approach differed from Brown and Robertson in several ways: First, Brown and Robertson (2007) excluded participants who exceeded a threshold for explicit sequence knowledge awareness, whereas we did not eliminate participants based on their level of sequence awareness (measured using the PDP). Rather, we included awareness as a factor in our statistical model. Our rationale for this deviation was that we did not want to introduce subjectivity into our analysis based on our chosen threshold for awareness, which may differ between age groups. Rather, we included awareness scores from the PDP as a factor in our analysis to isolate variance caused by sequence awareness from the potential interfering effect of episodic memory formation on procedural skill consolidation. Second, as opposed to using an ANOVA, we used a linear mixed-effect model analysis to isolate intersubject variability in scores from our fixed effects of interest. Our full analysis included TIME (categorical variable: P^1^ vs. P^2^), AGE (categorical variable: Young vs. Old), SUBGROUP (categorical variable: learning vs. control), AWARENESS (continuous variable: [-1, 1]), and their interactions as fixed factors, and subject intercept as a random effect. Our dependent variable was procedural skill performance. We expected a significant interaction between TIME, AGE, and SUBGROUP, indicating that memory interference was significantly different for older adults compared to younger adults. We planned to unpack the interaction by running separate linear mixed-effect model analyses for younger and older adults that included TIME, SUBGROUP, AWARENESS, and their interactions as fixed factors, and subject intercept as a random factor.

### Experiment 2 (P→E interference)

De-identified data (CSV files) and code (R markdown file; R-Studio version 4.2.3) needed to reproduce analyses for Experiment 2 are available for download at OpenICPSR.org (https://www.openicpsr.org/openicpsr/project/207361/version/V3/view). The experiment was pre-registered on aspredicted.org before conducting our analysis (#141678). We predicted that P→E memory interference would be significantly greater for older adults compared to younger adults

### Participants

Data collection for this experiment took place during the same time period as Experiment 1. Additionally, we recruited participants from the same area and the same inclusion and exclusion criteria were used. Our sample included 40 participants, consisting of 20 cognitively unimpaired young adults (22.9±3.28 years old) and 20 healthy older adults (65.6±6.97 years old).

Participants provided their informed consent and privacy rights of all participants were observed by all experiments. The Institutional Review Board at UT Austin approved all procedures on March 19^th^, 2021 (STUDY #00000860) and all participants were compensated for their participation. Brown and Robertson (2007) showed that procedural memory interferes with the consolidation of episodic memories in healthy younger adults. While the SRTT control group experienced virtually no change in episodic memory performance between laboratory visits (M = 0.0, SEM = 0.4), the SRTT learning group experienced offline losses (M = -1.6, SEM = 0.3).

This translates to an effect size of 1.43 (Cohen’s *d*). Assuming an alpha of 0.05 and power equal to 0.85, we would have an 85.6% chance of observing a significant memory interference effect in the younger adult age group with a sample size of 10 participants per sub-group (learning and control) using an unpaired, two-tailed t-test. Thus, all four groups included 10 participants.

### Procedures

Figure 1 (bottom) shows the experimental timeline for participants. In the morning, participants performed the word-list task (Ep^1^), as described above. Next, depending on the sub-group, participants either performed a procedural learning (SRTT with sequence) or control (SRTT without sequence) task. Finally, after a 6-12 h interval (average = 7 hours and 11 minutes ± 48), participants were retested on the word-list task during the afternoon session (Ep^2^). In general, the same tasks and stimuli were used in Experiment 2 as Experiment 1 save two differences: First, whereas both learning sub-groups (younger and older) performed an identical version of the SRTT as in Experiment 1, the control sub-groups performed a modified version of the SRTT where all cues were random, thus minimizing the potential for sequence learning. Second, we did not include the PDP to measure sequence awareness knowledge.

### Data analysis

Two participants in the Old-control group performed Experiment 1 prior to participating in this experiment. Both participants were part of the older adults-control group in Experiment 1 and 2 and experienced the word list described in Experiment, but not the embedded SRTT sequence. Though including these participants may have influenced our results, we included these participants for two reasons: 1) There was six months between experiments for each participant, which is a reasonable amount of time for memory of the list to fade, and 2) both participants did not perform at ceiling on the episodic memory task in Experiment 2.

Episodic memory performance was assessed during the morning session (Ep^1^) as recall after the line discrimination task and during the afternoon session (Ep^2^; Fig. 1**., bottom**). We calculated participants change in episodic memory performance as the difference between Ep^2^ and Ep^1^, where positive values indicate offline gains and negative values indicate offline losses. Procedural memory performance was calculated identically to Experiment 1 using the third SRTT block.

### Statistical analysis

Our full analysis included TIME (categorical variable: Ep^1^ vs. Ep^2^), AGE (categorical variable: Young vs. Old), SUBGROUP (categorical variable: learning vs. control), and their interactions as fixed factors, and subject intercept as a random effect. Our dependent variable was episodic memory performance. We expected to observe a significant interaction between TIME, AGE, and SUBGROUP, indicating that memory interference was significantly different for older adults compared to younger adults. We planned to unpack the latter interaction by running separate linear mixed-effect model analyses for younger and older adults that included TIME and SUBGROUP, and their interaction as fixed factors, and subject intercept as a random factor. In contrast to Experiment 1, participants did not perform the PDP, and we did not include awareness as a factor in our analysis. The rationale for this difference is that the time between the SRTT (morning) and awareness testing (afternoon) would be too long to reliably indicate sequence awareness. Additionally, the control group was not exposed to a sequence and thus, we would not be able to supply a full factorial set of awareness data to the analysis.

Our main results revealed neither strong evidence of memory interference in either group or a difference in the magnitude of P→E interference between age groups. These null results raise the possibility that our study was not sufficiently powered to observe these effects. As mentioned above, our power analysis was based on the twenty participants included in the second experiment of Brown and Robertson’s (2007) report, which revealed that we would have an 85.6% chance of observing a significant memory interference effect in younger adults with a sample size of 10 participants per sub-group (learning and control) using an unpaired, two-tailed t-test. However, while this power analysis indicates that we are powered to observe an interfering effect of procedural memory acquisition on episodic memory consolidation in younger adults, it does not necessarily indicate that we are powered to observe a significant interaction between memory interference and age group (younger vs. older). Because no studies to our knowledge have examined the effects of procedural memory acquisition on the wakeful consolidation of episodic memories in older adults, we performed post-hoc assessments of power from our garnered effect sizes to determine the number of participants necessary to observe a significant TIME x SUBGROUP (indicating memory interference in both groups) and SUBGROUP x TIME x AGE (indicating greater memory interference in one age group over the other) interaction using G power [33]. The rationale for this approach was to determine whether we should continue to collect more participants and retest our hypothesis with a lower alpha or whether doing so would require an impractical number of participants (e.g., n ≥ 1000 per group).

## Results

### Experiment 1 (E→P interference)

Figure 2A (left) shows P^1^ performance for all groups. We used an ANOVA to identify potential baseline differences between groups, but did not find any effect of SUBGROUP or AGE, nor a significant interaction. As expected, episodic memory performance after P^1^ was significantly greater in younger adults (14.6±0.50) compared to older adults (11.0±0.87; *F*(18) = 12.9, *p* < 0.005, *η_p_*^2^ = 0.42; Fig. 2B, **left**). Figure 2C (left) shows consolidation scores for all four sub-groups. We performed a linear mixed effects model analysis on procedural memory scores using AGE (younger vs. older), SUBGROUP (learning vs. control), TIME (morning vs. afternoon), AWARENESS, and their interactions as fixed factors, while separating these effects from subject-specific variability. Our analysis revealed a significant interaction between AGE and TIME (*t*(32) = 2.43, *p* < 0.05, *η_p_*^2^ = 0.002), indicating that the change in procedural memory performance between P^1^ and P^2^ was significantly lower for older adults compared to younger adults, regardless of sub-group. We also observed a significant SUBGROUP x TIME interaction (*t*(32) = 2.93, *p* < 0.01, *η_p_*^2^ = 0.03), indicating that the decrease in procedural memory performance between P^1^ and P^2^ was larger for the learning groups compared to the control groups, regardless of age group. Finally, we identified a significant AGE x SUBGROUP x TIME interaction, suggesting that the SUBGROUP x TIME interaction was stronger in older adults compared to younger adults (*t*(32) = -2.66, *p* < 0.05, *η_p_*^2^ = 0.057; Fig. 2C**, left**). No other effects or interactions reached significance. To unpack the three-way interaction between AGE, SUBGROUP, and TIME, we performed separate linear mixed effect model analyses for younger and older adults using SUBGROUP, TIME, AWARENESS, and their interactions as fixed factors, and subject intercept as a random factor. The analysis of younger adults revealed a significant interaction between TIME and AWARENESS (*t*(16) = -2.44, *p* < 0.05, *η_p_*^2^ = 0.16), indicating that awareness of the sequence was negatively associated with consolidation from P^1^ to P^2^ for younger adults, irrespective of sub-group. No other effects or interactions were significant (all *p*’s > 0.06). The analysis of older adults revealed a significant SUBGROUP x TIME interaction (*t*(16) = 2.55, *p* < 0.05, *η_p_*^2^ = 0.12), indicating a greater decrease in procedural memory performance from P^1^ to P^2^ for the learning group compared to the control group (see Fig. 2C**, left**). To determine whether episodic memory performance was related to skill consolidation, we regressed these variables against each other for both age groups. We did not identify significance for either group (Young: *r*(8) = 0.21, *ns*; Old: *r*(8) = 0.28, *ns*).

**Figure 2.**
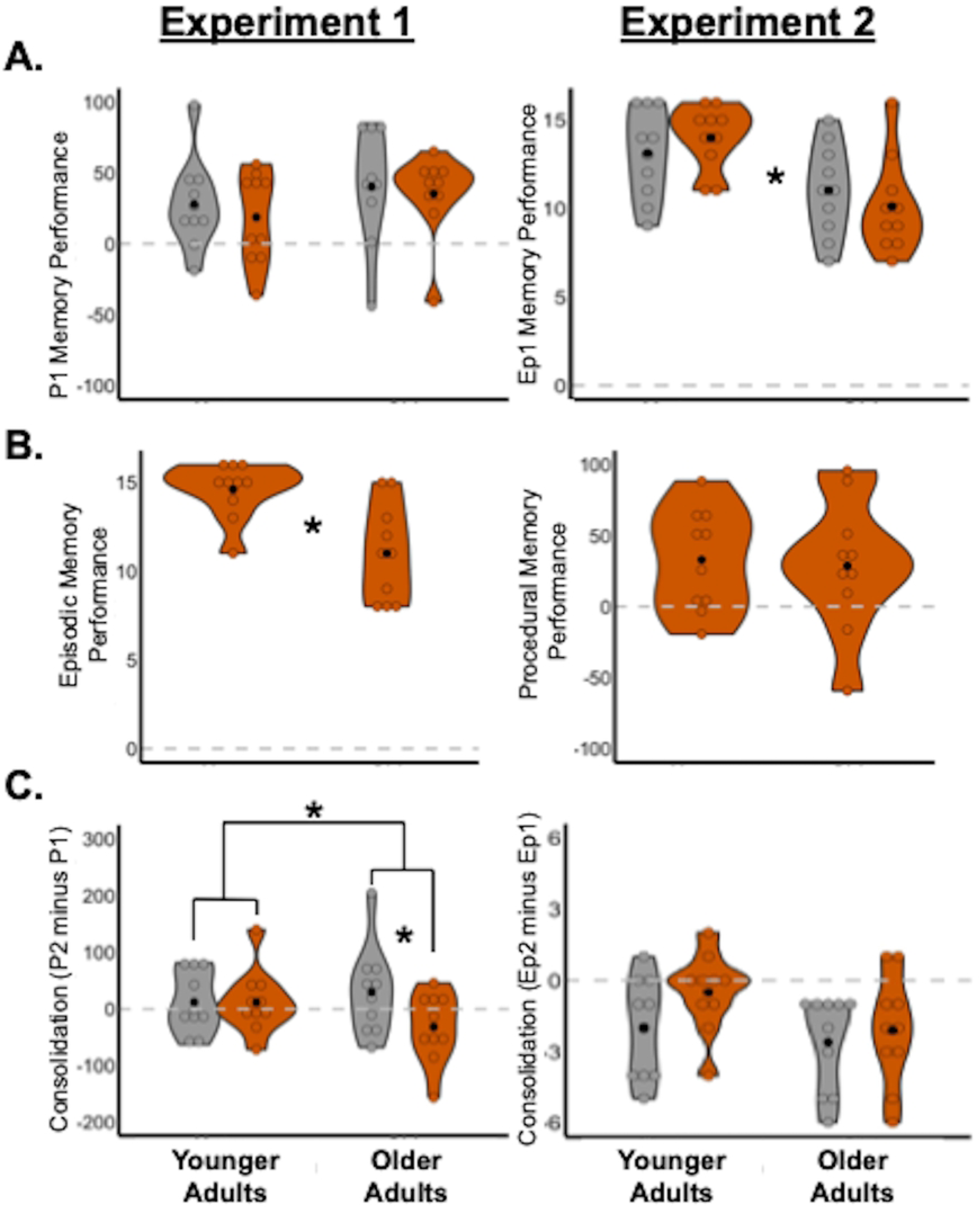
Experimental results. (A) Results for P^1^ (left) and Ep^1^ (right). (B) Results of word-list learning (left) and SRTT (right) learning tasks. (C) Consolidation results for Experiments 1 (left) and 2 (right), respectively. All violin plots show results for control (grey) and learning (orange) groups. Black dots represent mean and colored dots represent individual data points. *-*p*<0.05

### Experiment 2 (P→E interference)

Figure 2A (right) shows Ep^1^ performance for all groups. As expected, episodic memory performance was significantly greater in younger adults (13.6±0.49) compared to older adults (10.0±0.89; *F*(1,38) = 15.41, *p* < 0.005, *η_p_*^2^ = 0.29). Procedural memory performance after Ep^1^ was not significantly different between age groups; Fig. 2B**, right**). Figure 2C (right) shows consolidation scores for all four sub-groups. We performed a linear mixed effects model analysis on episodic memory scores using AGE (younger vs. older), SUBGROUP (learning vs. control), TIME (morning vs. afternoon), and their interactions as fixed factors, while separating these effects from subject-specific effects. Our analysis revealed a significant effect of TIME (*t*(36) = -3.28, *p* < 0.005, *η_p_*^2^ = 0.10), and a trend interaction between AGE and TIME (*t*(36) = 1.77, *p* = 0.09, *η_p_*^2^ = 0.01). The time effect indicates that episodic memory scores decreased from the morning to the afternoon, regardless of age or sub-group. The trending interaction suggests that older adults experienced a bigger decrease in episodic memory scores than younger adults. We found no strong evidence of procedural memory interference on episodic memory consolidation across participants of all ages (t(36) = -0.78, p = 0.44).

A determination of sample size using the partial eta squared score generated from an identical ANOVA (*η_p_*^2^ = 0.01) revealed that a sample of 384 participants (∼96 per group) would be necessary to observe a significant interaction between SUBGROUP and TIME (⍺ = 0.05, 1-β = 0.85). For the SUBGROUP, TIME, and AGEGROUP interaction (*η_p_*^2^ = 0.002), the same ANOVA revealed that a sample size of 6,143 participants (∼1,536 per group) would be necessary to observe a significant interaction between SUBGROUP, TIME, and AGEGROUP (⍺ = 0.05, 1-β = 0.85). The large number of participants required to see this effect suggests that it is unlikely that our null result was due to a lack of power.

## Discussion

Cross-memory interference occurs when the formation of episodic memory disrupts the consolidation of a procedural memory (E→P interference), and vice versa (P→E interference). Older adults experience reduced episodic memory, but relatively intact procedural memory [20,21], suggesting that one type of cross-memory interference may be exacerbated more than the other in this age group. To this end, we compared the magnitude of cross-memory interference between younger and older adults in two experiments. Our results show an asymmetrical increase in cross-memory interference: While E→P interference was greater in older adults, we found no statistical difference in P→E interference between age groups.

Collectively, our results indicate that aging does not uniformly affect cross-memory interference and suggest that this is asymmetric worsening of cross-memory interference is caused by the decline of the episodic memory system in older adults.

Our results provide a clue to determine how cross-memory interference occurs in younger and older adults. Based on our results, we can also hypothesize several mechanisms that cause increased E→P interference in older adults. Older adults experience reduced network specificity, which is the ratio between within and between-network functional connectivity. Specifically, they experience both decreased within-[34,35] and increased between-[36,37] network functional connectivity, and these neural changes are related to reduced cognitive performance [34,38–40]. Thus, the results we found could reasonably be caused by decreased episodic network efficiency, increased crosstalk between the episodic and procedural memory networks, or a combination of the two. For example, if E→P interference is caused by excitatory connections from the episodic to the procedural memory network, then degradation of the episodic memory network via aging may cause those inputs to become unregulated, increasing between-network functional connectivity and worsening E→P interference.

Another mechanism that can explain the increased E→P interference we observed in older adults is the process of dedifferentiation, which is loss of preferential network activation of the episodic and procedural memory networks: Dennis and colleagues (2011) found that while younger adults show preferential activation of the hippocampus and striatum during explicit and implicit learning, respectively, the same is not true for older adults [26]. Assuming that the striatum is performing double duty by supporting both episodic and procedural memory for older adults, it’s resources may be limited during procedural memory acquisition while episodic memory consolidation is also occurring, ultimately weakening procedural memory consolidation.

A final possibility is that greater episodic memory interference in older adults is caused by reduced posterior and increased frontal activation during memory tasks [41,42] compared to younger adults. Frontal brain regions are strongly associated with executive functions such as attention, inhibition, and cognitive control [43,44], which play crucial roles in memory encoding, retrieval, and interference resolution [45–47]. In contrast, posterior brain regions are particularly involved in sensory processing and perceptual encoding [48,49]. The shift from less posterior to more frontal activation patterns observed in older adults during episodic memory performance may contribute to greater interference in older adults by compromising cognitive control or attentional mechanisms dedicated to memory encoding and retrieval, forcing older adults to engage in compensatory strategies to acquire episodic memories, ultimately hurting procedural memory consolidation. Further research will be needed to adjudicate these possibilities.

In contrast to previous work [13], we did not identify strong evidence of cross-memory interference in younger adults. Some methodological differences between our studies might contribute to the disparate results. For example, differences in the duration of time interval between morning and follow-up visits could explain this difference. Brown and Robertson’s (2007) experiment required 12-hour intervals between morning and afternoon visits whereas we allowed participants to return between 6-12 hours after the morning visit. More specifically, on average, our participants took ∼6-7 hours between the beginning of the morning session and the start of the afternoon session in both experiments. As a result, less time between sessions could have caused less cross-memory interference. However, we did not find a relationship between interval time and memory interference in either of the learning groups across the two experiments. Differences in participant characteristics between experiments, such as age, could also affect susceptibility to interference and explain the difference in results. For example, participants in Brown and Robertson’s (2007) study were younger (21.1 ± 0.3) compared to the participants in our younger groups for Experiments 1 (24.35 ± 3.93) and 2 (25.70 ± 0.73). The age difference in younger adults may be associated with susceptibility to interference, although we did not find evidence for this relationship amongst younger adults. Another difference between the prior report and our experiments is that we added a perceptual distraction task to discourage rehearsal between word list learning and recall in the episodic memory task. The perceptual distraction task could have reduced episodic memory recall and consequently minimized the possibility for interference. However, younger adults had significantly stronger episodic memory recall than older adults and yet we only found significant episodic memory interference on procedural skill consolidation only in older adults. Thus, it is unlikely that the perceptual distraction task alone explains the difference in results between studies. Finally, we included the PDP [30] to assess conscious control of sequence awareness at the end of the afternoon visits, whereas Brown and Robertson (2007) “removed participants who were able to recall four or more items from the 12-item sequence”[13]. As individuals become proficient in executing a sequence during the SRTT task, they also develop the capability to verbally explain the sequence, which can affect procedural skill consolidation [50]. To overcome this, we statistically isolated variance associated with awareness from our fixed effects of interest. Nevertheless, any or all of these factors could have contributed to our null findings for younger adults.

Our study had several strengths. First, we used a linear mixed effects model to separate inter-individual variability in memory performance from our fixed effects of interest. Second, we used awareness scores from the PDP as a factor in our Experiment 1 analysis to separate out variance caused by conscious sequence control instead of removing participants from our analysis. Third, we included a perceptual distracting task to prevent age-related differences in the rehearsal of memories during the episodic memory tasks. The most significant limitation of our study is the shorter time interval between morning and afternoon sessions compared to Brown and Robertson (2007). A shorter interval may produce weaker interference, although six hours was sufficient to observe this effect in a conceptual replication of Brown and Robertson’s (2007) study [14]. Another limitation was our sample sizes. Larger sample sizes allow for more robust statistical comparisons and increase the reliability of results. However, our power analysis indicated that ten participants per group should yield sufficient power to reveal memory interference in the younger group and our *post hoc* analyses showed that thousands of participants would be required to power a study to observe age-related memory interference differences in Experiment 2. The present study also focused on only behavioral aspects of memory interference in younger and older adults. More work will be necessary to determine the mechanistic underpinnings causing greater memory interference in older adults. Despite these limitations, the strengths of our study contribute to a more comprehensive understanding of episodic-procedural memory dynamics in older adults, and we provide several possible mechanisms to explain why older adults experience greater E→P interference. Studies using functional task-based neuroimaging and larger sample sizes will provide a more comprehensive understanding of how episodic memory interferes with wakeful procedural consolidation in both age groups.

## Conclusions

In sum, we compared the magnitude of the interfering effects of episodic memory acquisition on procedural skill consolidation between healthy younger (18-40 years old) and older (≥ 55 years old) adults. Our findings demonstrate that episodic memory interference on procedural skill consolidation is significantly greater in older adults compared to younger adults, indicating asymmetrical worsening of cross-memory interference and suggesting that E→P interference is related to episodic memory decline in older adults.

